# Identification of C270 as a novel site for allosteric modulators of SARS-CoV-2 papain-like protease

**DOI:** 10.1101/2022.03.30.486313

**Authors:** Hangchen Hu, Qian Wang, Haixia Su, Qiang Shao, Wenfeng Zhao, Guofeng Chen, Minjun Li, Yechun Xu

## Abstract

The papain-like protease (PL^pro^) in coronavirus is one of key cysteine proteases responsible for the proteolytic processing of viral polyproteins, and plays an important role in dysregulation of host immune response. PL^pro^ is a promising therapeutic target with a major challenge in inhibitor design due to the restricted S1/S2 sites for two consecutive glycine of substrates. Here we reported the discovery of two activators of the SARS-CoV-2 PL^pro^ from a biochemical screening, and the identification of the unique residue, C270, as an allosteric and covalent regulation site for the activators. This site was also specifically modified by glutathione oxidized, resulting in the S-glutathionylation and activation of the protease. Furthermore, one compound was found to allosterically inhibit the protease by covalent binding to this crucial site. Together, these results elucidated an unrevealed molecular mechanism for allosteric modulation of the protease’s activity, and provided a new strategy for discovery of allosteric inhibitors of the SARS-CoV-2 PL^pro^.

## Introduction

Coronaviruses (CoVs) are highly diverse, enveloped, positive-sense, single-stranded RNA viruses, which are responsible for a wide range of diseases in animals as well as humans^[1-2]^. The COVID-19 pandemic is caused by a new human CoV, severe acute respiratory syndrome coronavirus 2 (SARS-CoV-2) that appears to be more readily transmitted from human to human^[3-4]^. Besides SARS-CoV-2, human infections with zoonotic CoVs including SARS in 2003 and Middle East respiratory syndrome (MERS) in 2012 caused severe respiratory diseases with high morbidity and mortality, posing a significant threat to public health across the world^[5-6]^. SARS-CoV-2 was 96% identical at the whole-genome level to a bat coronavirus and shares 79.6% sequence identity to SARS-CoV^[7]^. The ongoing pandemic has caused unbearable number of infections and deaths worldwide which are still increasing, rendering it the most horrible pandemic since the 1918 Spanish flu^[8]^. It also marks the third introduction of a highly pathogenic CoV into the human population in the 21th century^[9]^.

CoVs share key genomic elements that provide promising therapeutic targets^[10]^. A chymotrypsin-like cysteine protease, also called 3C-like protease (3CL^pro^) or main protease (M^pro^)^[11]^, together with a papain-like protease (PL^pro^)^[12]^, is required to process two viral polyproteins into mature nonstructural proteins which are essential for viral transcription and replication. Besides, PL^pro^ also recognizes and cleaves the C-terminal LxGG sequence of ubiquitin (Ub) and ubiquitin-like proteins (UbL) such as interferon-stimulated gene product 15, acting as a deubiquitinase (DUB) to remove Ub or UbL from host proteins^[13-17]^. Antiviral immunity includes broad posttranslational modification by Ub and UbL which regulate host protein cellular localization and stability. The DUB activity of PL^pro^ is supposed to cause dysregulation of host immune response against viral infection^[18-22]^. Thus, targeting PL^pro^ is an attractive strategy to both inhibit viral replication and prevent disruption of the host immune response against viral infection.

Significant progress has been made in development of inhibitors targeting the SARS-CoV-2 3CL^pro [23-27]^. Discovery of PF-00835231 as a covalent active-site-directed inhibitor of the SARS-CoV 3CL^pro^ in 2003 is conducive to rapidly develop inhibitors of the SARS-CoV-2 3CL^pro[24-25, 28-29]^. Currently, the most advanced inhibitor, nirmatrelvir (PF-07321332), in combination with ritonavir has been approved for the treatment of COVID-19^[28]^. In comparison, PL^pro^ of SARS-CoV and SARS-CoV-2 represents a more challenging target as very few potent inhibitors of this protease have been reported with validated in vitro as well as in vivo efficacy. A key issue from the standpoint of inhibitor design is the barrier derived from the restricted binding sites (S1/S2) for recognizing two consecutive glycine (P1/P2) in substrates of PL^pro [30-32]^. Despite several high-throughput screening campaigns have been performed^[13, 31, 33-36]^, GRL0617 which was originally identified as an inhibitor of the SARS-CoV PL^pro[37]^, remains one of the most potent PL^pro^ inhibitors and the major starting point for optimization^[17, 30, 38-41]^. Comprehensive understanding of structure-activity relationships of the protease and exploration of more ligand binding sites in addition to the catalytic (orthosteric) site is thus required for new potent inhibitor design.

In this work, we performed a high-throughput screening of compounds including FDA-approved drugs and candidates in clinical trials using an enzymatic assay. Though we embarked on our search for new inhibitors of the SARS-CoV-2 PL^pro^, we unexpectedly found two disulfide-containing activators which subsequently enabled us to identify a novel binding site on the SARS-CoV-2 PL^pro^ and elucidated an unrevealed allosteric and covalent binding mechanism to regulate activity of the protease. Inspired by this discovery, an endogenous activator and the first allosteric inhibitor were also identified to covalently modify the protease through this site. These findings thereby gain the mechanistic insights into activity regulation of the SARS-CoV-2 PL^pro^ and find a vital allosteric site for the development of novel inhibitors against SARS-CoV-2.

## Results and discussion

### Discovery of two activators of the SARS-CoV-2 PL^pro^ with novel mechanism

To measure the proteolytic activity of the recombinant SARS-CoV-2 PL^pro^, we carried out an enzymatic assay with a short fluorogeneic substrate, RLRGG-AMC, which is labelled with 7-amino-4-methylcoumarin (AMC)^[42]^. The hydrolysis of this substrate by the protease to release the AMC group results in a substantial enhancement of the fluorescent intensity. With this assay, FDA-approved drugs and candidate compounds in clinical trials (100 μM final concentration) were tested against the SARS-CoV-2 PL^pro^ (50 nM final concentration) in order to screen inhibitors of the protease. Intriguingly, a number of activators which increase the proteolytic activity of the protease were resulted from the screening. Among them, two compounds both containing a disulfide bond, dimesna and pyritinol, led to more than 50% activation of the SARS-CoV-2 PL^pro^ (Figure 1a). Moreover, both compounds induced significant upregulation of the protease’s activity in a dose-dependent manner (Figure S1). We subsequently determined the values of half maximal effective concentration (EC_50_) and maximal effect (E_max_) representing activation potency and efficacy, respectively, of the two activators. Dimesna, an uroprotective agent, activated the SARS-CoV-2 PL^pro^ with an EC_50_ value of 1046 μM and an E_max_ value of 300.0% (Figure 1a and Figure S1). The other activator, pyritinol which is an analog of vitamin B6 for the treatment of cognitive disorders, activated the protease with an EC_50_ value of 18 μM and an E_max_ value of 226.5% (Figure 1a and Figure S1). The effect of these two compounds on Michaelis constant (*K*_m_) and maximum reaction rate (*V*_max_) of the fluorogeneic substrate hydrolyzed by the SARS-CoV-2 PL^pro^ was also determined (Table 1). Both activators rarely have an influence on *K*_m_ but increased the *V*_max_ values, indicating that the activators enhance the reaction rate rather than binding affinity of the substrate with the protease. Taken together, dimesna and pyritinol were found to be the allosteric activator of the SARS-CoV-2 PL^pro^ for the first time.

**Table 1.**
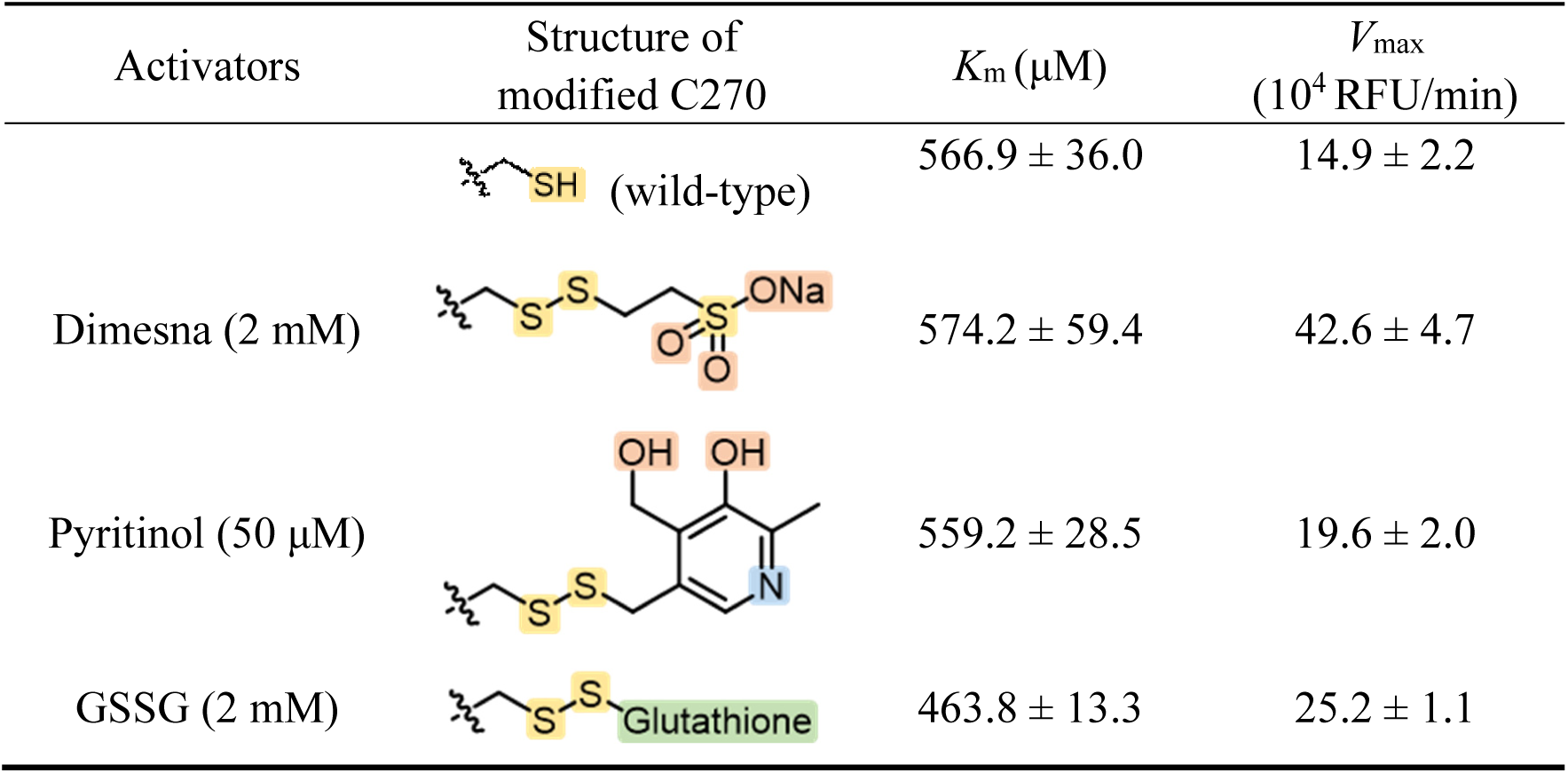
*K*_m_ and *V*_max_ values of the fluorogeneic substrate (RLRGG-AMC) hydrolyzed by the wild-type SARS-CoV-2 PL^pro^ in the absent or present of activators.

| Activators | Structure of modified C270 | $K_m$ ($\mu M$) | $V_{max}$ ($10^4$ RFU/min) |
| --- | --- | --- | --- |
| | (wild-type) | $566.9 \pm 36.0$ | $14.9 \pm 2.2$ |
| Dimesna (2 mM) | | $574.2 \pm 59.4$ | $42.6 \pm 4.7$ |
| Pyritinol (50 $\mu M$) | | $559.2 \pm 28.5$ | $19.6 \pm 2.0$ |
| GSSG (2 mM) | | $463.8 \pm 13.3$ | $25.2 \pm 1.1$ |

**Figure 1.**
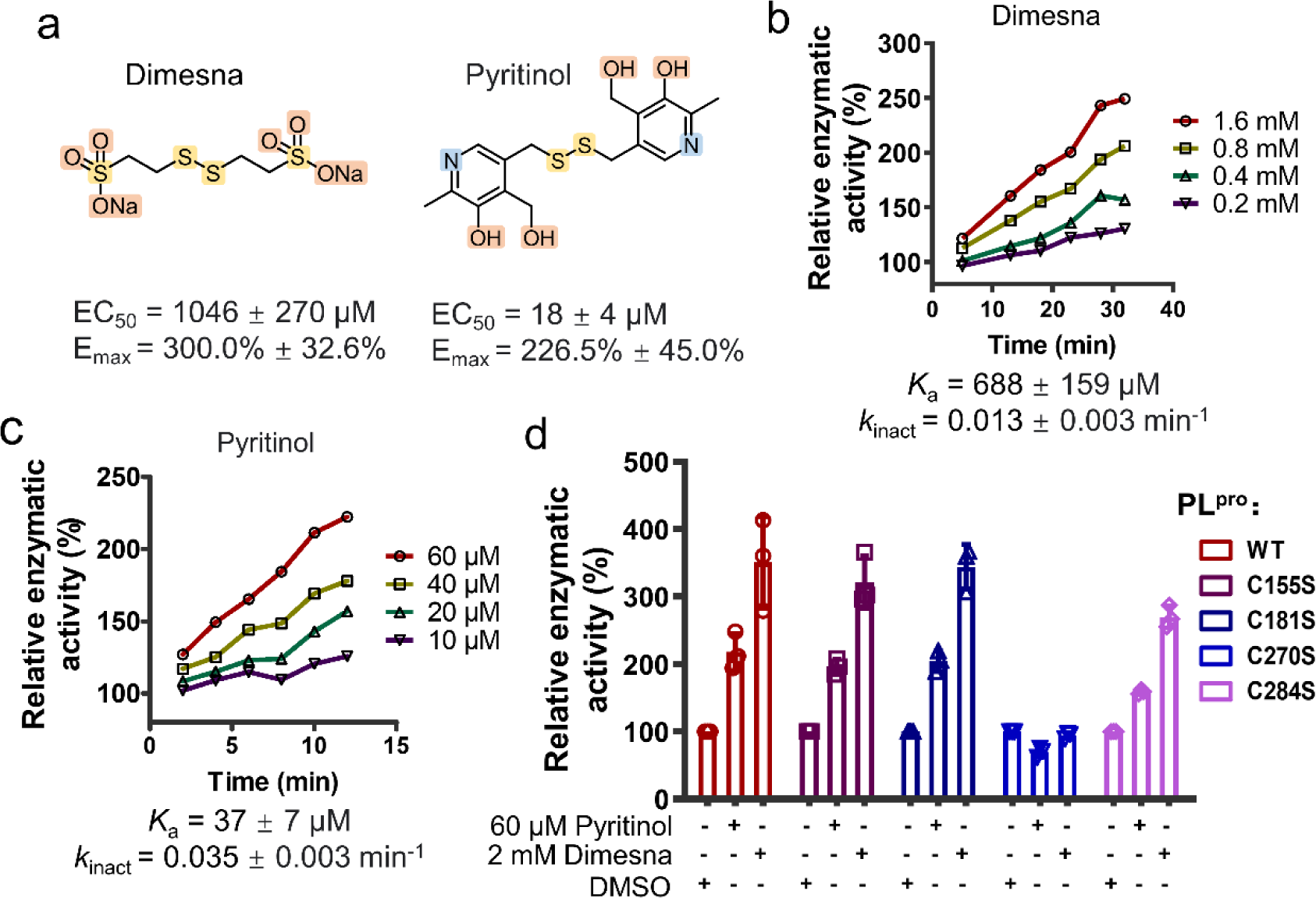
Dimesna and pyritinol upregulated the enzymatic activity of SARS-CoV-2 PL^pro^ by covalently binding to C270. (a) Chemical structures, activation potency and efficacy (EC_50_ and E_max_) of dimesna and pyritinol. (b-c) Time-dependent activation of the SARS-CoV-2 PL^pro^ by dimesna (b) and pyritinol (c) at various concentrations, and the calculated covalent binding kinetic parameters. (d) Activation effect of dimesna and pyritinol on the wild-type (WT) SARS-CoV-2 PL^pro^ and its four variants, C155S, C181S, C270S, and C284S.

In addition, two compounds showed time-dependent activation of the SARS-CoV-2 PL^pro^ (Figure 1b and Figure 1c), which is in line with the model of covalent modulation. Covalent ligand binding involves a two-step process: an initial reversible binding event followed by formation of the covalent bond, which is characterized by the binding affinity (*K*_a_ for activators, *K*_i_ for inhibitors) and the rate constant of covalent bond formation (*k*_inact_), respectively. To obtain kinetic parameters of the activators binding to the SARS-CoV-2 PL^pro^, various concentrations of the compound were incubated with the protease at a final concentration of 50 nM for several indicated time, and activities of the protease were measured after incubation (Figure 1b and Figure 1c). The result showed that two activators had comparable *k*_inact_ values, which might be ascribed to the same reaction group, the disulfide bond. However, pyritinol had a much lower *K*_a_ value than that of dimesna, which suggests more noncovalent interactions formed between pyritinol and the protease. Therefore, dimesna and pyritinol are two covalent activators of the SARS-CoV-2 PL^pro^ with the same reaction group but different amount of noncovalent contacts.

Considering that compounds containing a disulfide bond can act as covalent probes to characterize cysteine at allosteric sites^[43-48]^, we predicted that the activity modulation of the SARS-CoV-2 PL^pro^ by dimesna and pyritinol might be resulted from the modification of cysteine residue(s) of the protease through an action of disulfide exchange. To test this prediction, four vulnerable cysteine on the surface of the protease, C155, C181, C270, and C284, were individually mutated to serine. The resulting variants of the recombinant SARS-CoV-2 PL^pro^ C155S, C181S, C270S, and C284S, were expressed and purified for the proteolytic activity measurement without or with addition of the activator (Figure 1d). It was revealed that the enzymatic activity of all the variants except C270S were upregulated by the two activators, demonstrating that C270 of the SARS-CoV-2 PL^pro^ was the modification site for the activators. It has been reported that C270 has the lowest p*K*a among all the cysteine of the SARS-CoV-2 PL^pro^ except for the catalytic cysteine (C111)^[49]^, which means that C270 has a high reactivity and tendency for covalent modification. Though the catalytic C111 has the highest reactivity, it is not easy to be modified by two activators as it is located inside the protease and still works for catalytic hydrolysis of the substrate. Consequently, the enzymatic activity examination of the protease with single mutation of several surface cysteine reveals that C270 was modified by the two activators and this led to the activity upregulation of the SARS-CoV-2 PL^pro^, which is consistent with our prediction.

### Glutathione oxidized (GSSG) activated the SARS-CoV-2 PL^pro^ through C270 modification

The confirmation that C270 was the modulate site for two disulfide-containing compounds to active the SARS-CoV-2 PL^pro^ turned our attention to an endogenous molecule, GSSG, which also contains a disulfide bond and has potential to modify the protease in this manner. To support this speculation, GSSG was tested and showed to upregulate the protease activity as anticipated (Figure 2a). In addition, such activation was eliminated by the introduce of the C270S mutant (Figure 2a), which confirmed that the activity regulation was achieved through the modification of C270. The effect of GSSG on the *K*_m_ and *V*_max_ values of the SARS-CoV-2 PL^pro^ was investigated (Table 1). Similar to the two activators mentioned above, GSSG hardly had an influence on the *K*_m_ but increased the *V*_max_ value (25.2 vs. 14.9). GSSG was also found to exert the time-dependent activation on the SARS-CoV-2 PL^pro^, which is in line with an irreversible binding mode (Figure 2b). Kinetic parameters of GSSG binding to the SARS-CoV-2 PL^pro^ were calculated based on the measurement of the protease activity after incubation with various concentrations of GSSG for indicated time. As a result, GSSG resulted in *k*_inact_ and *K*_a_ values similar to those of dimesna. These data demonstrated that GSSG upregulates the protease’s activity in the same manner utilized by dimesna and pyritinol. To further assess the covalent modification of C270 by GSSG, a commercial antibody which is able to detect the S-glutathionylated protein was used for immunoblotting test (Figure 2c). Both the wild-type and C270S mutant of SARS-CoV-2 PL^pro^ were incubated with GSSG, and the resulting complexes were then detected by the antibody. The result showed that only the wild-type protease incubated with GSSG and then denatured under the unreduced condition showed a clear band in the immunoblotting test (Figure 2c), providing the evidence for the S-glutathionylation of C270 by GSSG. Moreover, this band was diminished under the reduced condition, which suggests the covalent linkage of GSSG to C270 of the protease through the disulfide exchange. In contrast, the C270S mutant of SARS-CoV-2 PL^pro^ treated with GSSG or not didn’t show any band (Figure 2c). It is thus concluded that the modification of GSSG on the protease mainly occurred at C270 rather than other cysteine.

**Figure 2.**
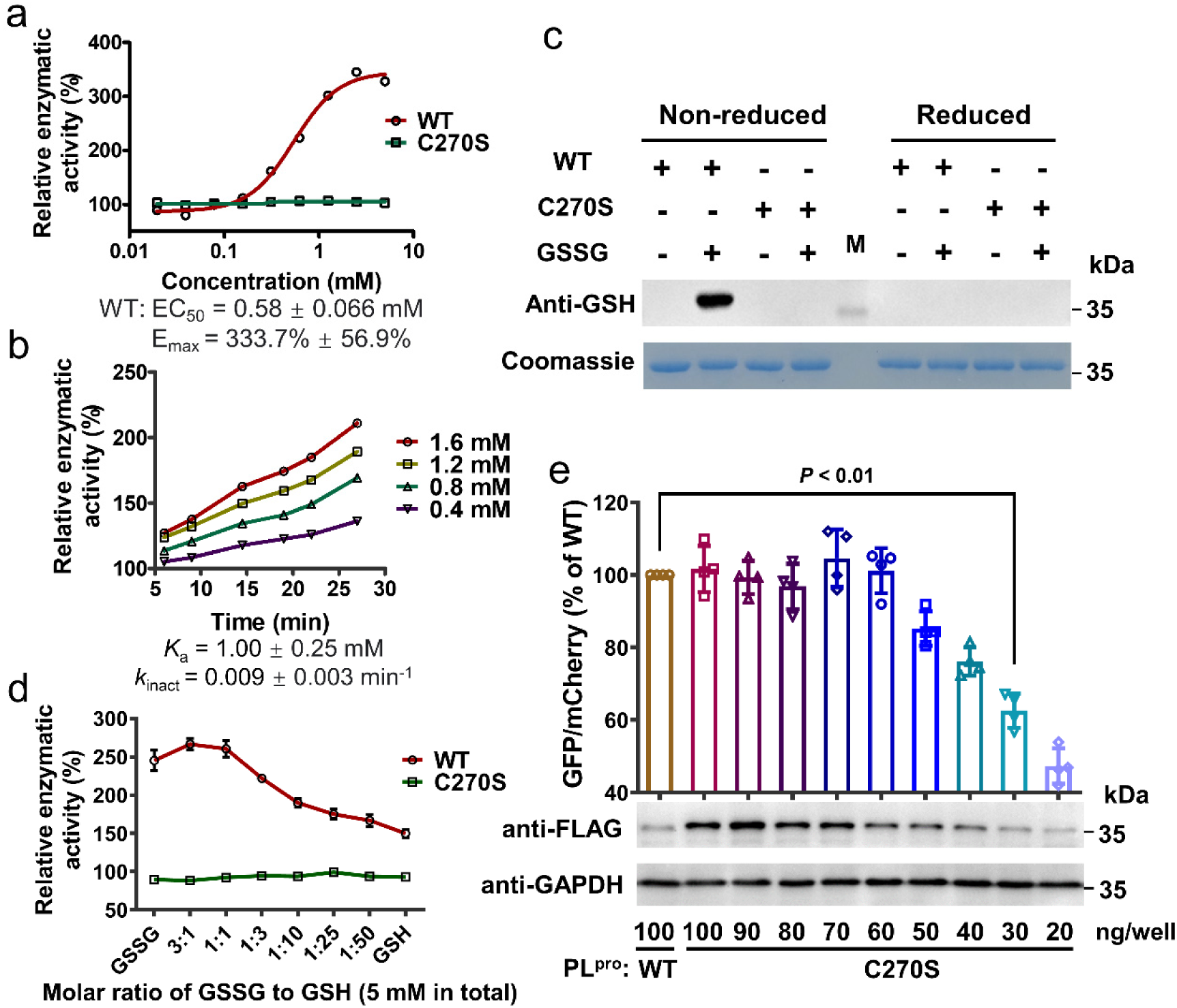
GSSG modified C270 of the SARS-CoV-2 PL^pro^ to enhance the enzymatic activity of the protease. (a) Representative profiles for the activation of the wild-type (red) and C270S mutant (green) of SARS-CoV-2 PL^pro^ by GSSG. (b) Time-dependent activation of the SARS-CoV-2 PL^pro^ by GSSG at various concentrations, and the calculated covalent binding kinetic parameters. (c) Immunoblotting of the S-glutathionylation of the wild-type (WT) or C270S mutant of SARS-CoV-2 PL^pro^ with indicated treatments. (d) The activity of the WT or C270S mutant of SARS-CoV-2 PL^pro^ after treatment with various ratio of GSSG to GSH (5 mM for total glutathione). (e) Intracellular activity of the WT or C270S variant of SARS-CoV-2 PL^pro^ after transfection with different amount of plasmids. Immunoblotting evaluated the protein expression level of the WT and mutated protease in cells.

It is worth noting that the physiological concentration of total glutathione in cells ranges from 1 to 10 mM, while the ratio of reduced to oxidized glutathione (GSH/GSSG) varies depending on the different condition and organelle of cells^[50-51]^. Accordingly, we tested different ratios of GSH/GSSG at 5 mM for a total concentration of glutathione. The enzymatic activity of the wild-type SARS-CoV-2 PL^pro^ was upregulated at each ratio while it wasn’t changed for the C270S variant (Figure 2d). This result indicates that the proteolytic activity of SARS-CoV-2 PL^pro^ could be modulated by GSH/GSSG under the cellular conditions. In this context, a PL-FlipGFP assay ^[31]^ was performed to determine the intracellular activity of wild-type SARS-CoV-2 PL^pro^ and the C270S variant (Figure S2). In this assay, a fluorogenic GFP containing a cleavage site (LRGGAPTK) by the SARS-CoV-2 PL^pro^, namely PL-FlipGFP, and a red fluorescent protein (mCherry) were constructed together as the substrate of the protease and the internal control, respectively. The fluorescence of PL-FlipGFP was produced after the cleavage by the protease (Figure S2, see Supporting Information for detailed mechanism). Consequently, the ratio of GFP/mCherry fluorescent signal was used to represent the amount of hydrolyzed product by the protease in cells. The expression level of the FLAG-tagged protease was also detected by immunoblotting.

The result showed that the GFP/mCherry ratio of the wild-type and the C270S variant were nearly the same (100% vs. 101.6%) when transfected with the same amount of plasmids (Figure 2e and Figure S3). However, at this condition, the expression level of C270S variant was much higher than that of the wild-type. Since the GFP/mCherry ratio implied that the amount of hydrolyzed product resulting from two cases was similar, the higher expression level of C270S variant over the wild-type reflected the lower activity of C270S variant over the wild-type of SARS-CoV-2 PL^pro^ in cells. Nevertheless, this situation might also be caused by the insufficient substrates. To exclude this possibility, the amount of plasmid expressing the C270S variant was reduced to achieve the expression level similar to that of the wild-type. Consequently, the GFP/mCherry ratio of the wild-type was significant higher than that of the C270S variant (100% vs. 62.5%), demonstrating that the more hydrolyzed products were produced by the wild-type when the amount of expressed protein for two proteases was similar (Figure 2e). Taken together, these data revealed the higher activity of the wild-type over the C270S variant of SARS-CoV-2 PL^pro^ in cells which contains millimolar glutathione with different ratios of GSH/GSSG. It is thereby speculated that the performance of the intracellular proteolytic activity of SARS-CoV-2 PL^pro^ on the viral polyproteins might be activated by the endogenous glutathione.

### Impact of C270 mutants on the enzymatic activity of SARS-CoV-2 PL^pro^

The covalent modification of C270 by dimesna, pyritinol and GSSG leads to the activity upregulation of SARS-CoV-2 PL^pro^ while the C270S mutant eliminates such an activation action, underscoring the importance of C270 for modulation of the enzymatic activity of the protease. This raises the question whether the modification of C270 can inhibit the activity of the protease. To answer it and to aid the future efforts in developing novel inhibitors targeting this site, we first explored the influence of different mutants of C270 on the *K*_m_ and *V*_max_ of the protease. Considering different sizes, polarities and charge states of the side chain, eleven mutagenesis including C270S, C270V, C270L, C270E, C270Q, C270M, C270F, C270Y, C270H, C270K, and C270R, were carried out. Table 2 showed that all the mutants didn’t have significant influence on the *K*_m_ in comparison to the *K*_m_ value of the wild-type enzyme. As for *V*_max_, the mutagenesis such as C270S, C270V and C270L didn’t make a difference to the wild-type. The C270E mutant with the negative charge was originally designed to resemble the modification of C270 by dimesna, but its *V*_max_ value is much less than the one resulted from the addition of dimesna (11.4 vs. 42.6) and is even less than that of the wild-type (11.4 vs. 14.9). Additionally, the *V*_max_ values of C270M (21.6), C270F (26.0) and C270Y (26.5) mutants increased in comparison to that of the wild-type (14.9), which may be ascribed to relatively large and electron-rich side chains of methionine, phenylalanine and tyrosine. In contrast, C270H, C270K and C270R mutants which all contain a positive-charged side chain resulted in values of *V*_max_ smaller than that of the wild-type (9.3, 6.0, 4.7 vs. 14.9), suggesting that those mutagenesis caused an inhibition effect on the protease. Together, these results revealed that the substitution of C270 with amino acids containing large side chains to mimic the modification of C270 by compounds would affect the enzymatic activity of the protease, although detailed explanations for these results remain to be determined. It manifests again that C270 is a crucial allosteric site to modulate the protease’s activity, and suggests that the covalent modification by the positive-charged molecule may exhibit an inhibition effect on the protease.

**Table 2.**
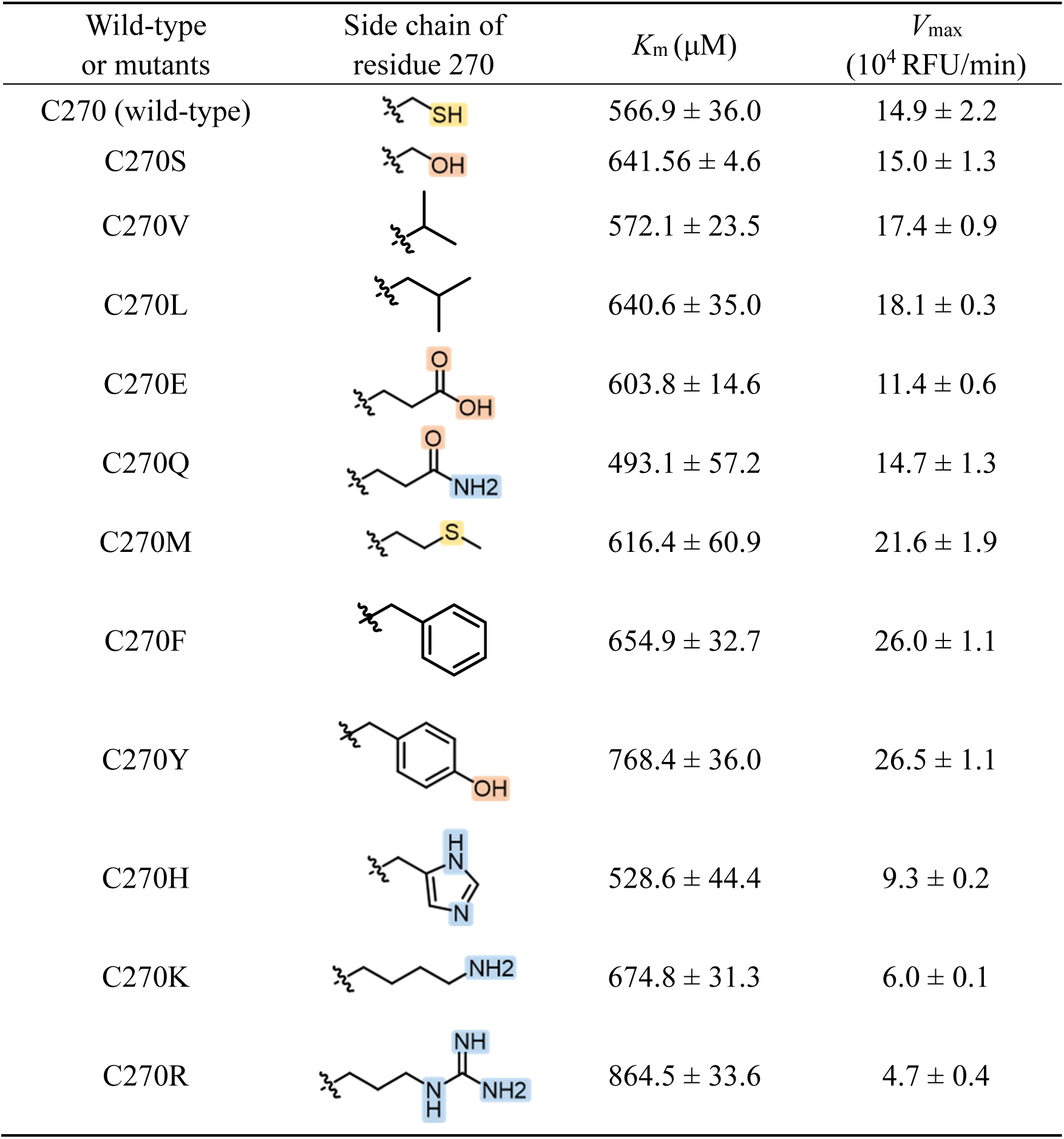
*K*_m_ and *V*_max_ values of the SARS-CoV-2 PL^pro^ with different mutations of C270.

| Wild-type or mutants | Side chain of residue 270 | $K_m$ (μM) | $V_{max}$ ($10^4$ RFU/min) |
| --- | --- | --- | --- |
| C270 (wild-type) | | $566.9 \pm 36.0$ | $14.9 \pm 2.2$ |
| C270S | | $641.56 \pm 4.6$ | $15.0 \pm 1.3$ |
| C270V | | $572.1 \pm 23.5$ | $17.4 \pm 0.9$ |
| C270L | | $640.6 \pm 35.0$ | $18.1 \pm 0.3$ |
| C270E | | $603.8 \pm 14.6$ | $11.4 \pm 0.6$ |
| C270Q | | $493.1 \pm 57.2$ | $14.7 \pm 1.3$ |
| C270M | | $616.4 \pm 60.9$ | $21.6 \pm 1.9$ |
| C270F | | $654.9 \pm 32.7$ | $26.0 \pm 1.1$ |
| C270Y | | $768.4 \pm 36.0$ | $26.5 \pm 1.1$ |
| C270H | | $528.6 \pm 44.4$ | $9.3 \pm 0.2$ |
| C270K | | $674.8 \pm 31.3$ | $6.0 \pm 0.1$ |
| C270R | | $864.5 \pm 33.6$ | $4.7 \pm 0.4$ |

### Discovery of novel allosteric inhibitor of the SARS-CoV-2 PL^pro^

With the aid of all suggestions from the above results, bis[2-(N,N-dimethylamino)ethyl] disulfide (DMGA), which contains a disulfide bond and two positive-charged methylamine, was found to perfectly meet the requirement for the inhibitor (Figure 3a). As anticipated, DMGA exhibited a dose-dependent inhibition of the SARS-CoV-2 PL^pro^ and the resulting IC_50_ value was 9.4 μM (Figure 3b). Activity measurements of the wild-type and the C270S mutant of the protease incubated with DMGA revealed that the inhibitory potency of DMGA was caused by the covalent modification of C270 (Figure 3c). In addition, DMGA showed a dose-dependent reduce of the *V*_max_ but hardly affected the *K*_m_ except for the highest concentration (40 μM) at which the *K*_m_ value increased from 566.9 μM to 798.1 μM (Figure 3d). These results are in line with the model of non-competitive inhibition. Moreover, DMGA was found to time-dependent inhibit the SARS-CoV-2 PL^pro^, demonstrating the covalent binding mode utilized by DMGA too (Figure 3e). Kinetic parameters of DMGA binding with the SARS-CoV-2 PL^pro^ were calculated based on the activity measurements of the protease incubated with various concentrations of DMGA for indicated time. The resulting *k*_inact_ and *K*_i_ value is 0.034 min^-1^ and 41 μM, respectively, which are similar to the values of pyritinol (Figure 3e and Figure 1c). These data together indicated that DMGA allosterically inhibits the SARS-CoV-2 PL^pro^ through a specific covalent modification on C270, providing convincing evidence that C270 is a pivotal site for design of novel allosteric inhibitors of the protease.

**Figure 3.**
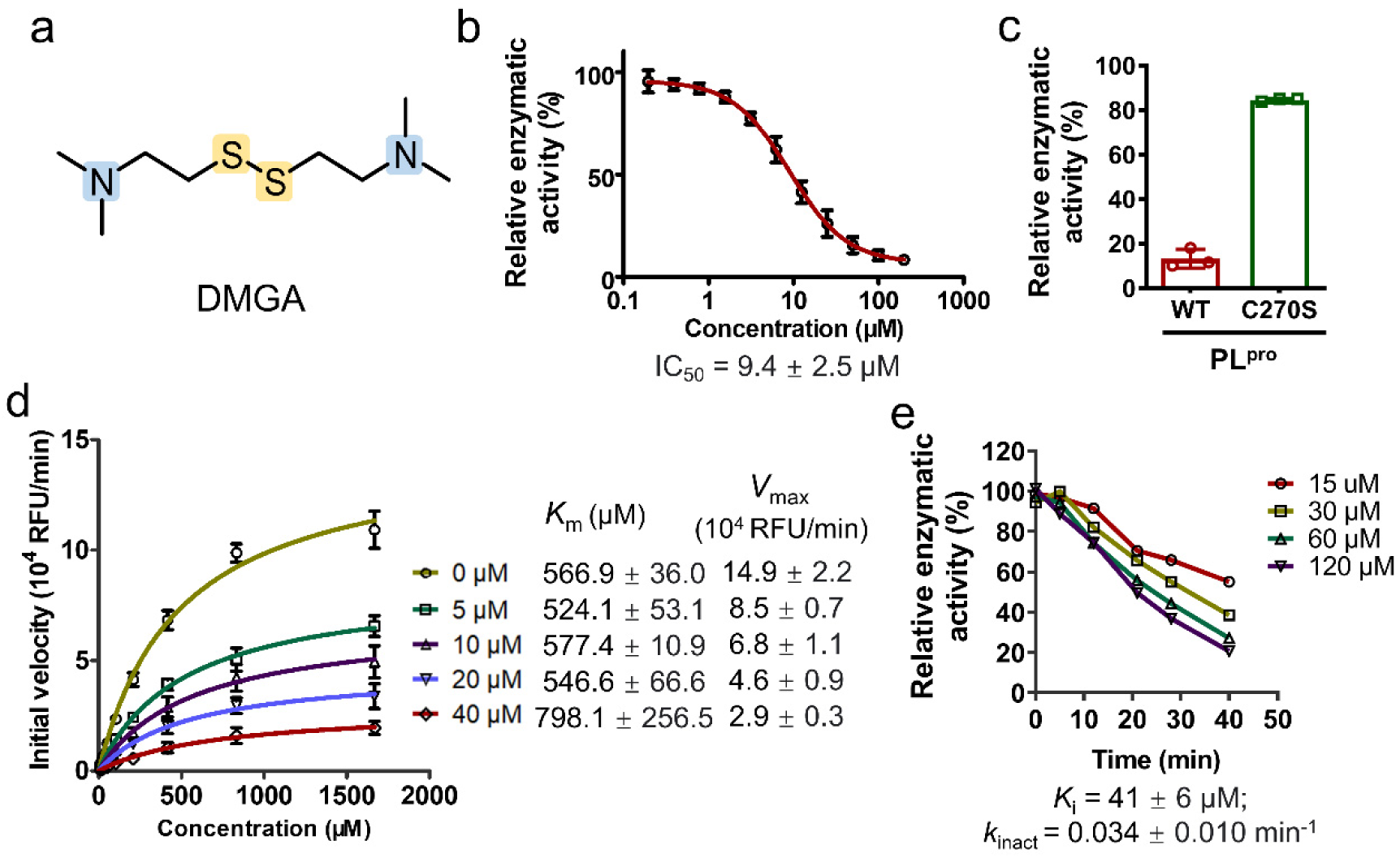
DMGA allosterically inhibits the SARS-CoV-2 PL^pro^ by covalently binding to C270. (a) Chemical structure of DMGA. (b) Representative profile for DMGA inhibiting the SARS-CoV-2 PL^pro^. (c) Different inhibitory rates of 50 μM DMGA on the wild-type and C270S mutant of SARS-CoV-2 PL^pro^. (d) The determination of *K*_m_ and *V*_max_ values of the substrate hydrolyzed by the SARS-CoV-2 PL^pro^ in the present of different concentrations (0-40 μM) of DMGA. (e) Time-dependent inhibition of the SARS-CoV-2 PL^pro^ by DMGA at various concentrations and indicated time was determined to calculate kinetic parameters of DMGA binding to and reacting with the protease.

It has been noted in Introduction that the discovery of PL^pro^ inhibitor is hampered by the restricted S1/S2 site for the binding of the tandem glycine in substrates. Such a space restriction also prevents the covalently binding of ligand to the catalytic C111. The discovery of C270 as an allosteric site thus provide an alternative route to inhibitor design. DMGA with single digit micromolar potency acts as a hit for further development of potent allosteric inhibitor.

In summary, we have investigated the activity modulation of the SARS-CoV-2 PL^pro^ by three activators, one inhibitor and eleven single-point mutants which consistently have an impact on the *V*_max_ but not the *K*_m_ of the protease by exerting the covalent modification on the surface C270, demonstrating the significance as well as feasibility of C270 working as an allosteric site for all the modulators. Noteworthy, the activators seen in the screening campaigns of inhibitors is ignored in general, while it is the fact that these activators frequently bind to an allosteric site which is potentially valuable for novel inhibitor design. The present study provides such a paradigm for the discovery of novel allosteric inhibitors of the SARS-CoV-2 PL^pro^ by utilizing the serendipitously discovered activators to identify the allosteric binding site.

To investigate the sequence conservation of C270 among PL^pro^, we performed a sequence alignment of pp1ab from seven human CoVs (Figure S4). It was revealed that C270 is unique to the SARS-CoV and SARS-CoV-2 PL^pro^, and instead valine was found in the other five CoVs’ PL^pro^ at the equivalent position. Considering that only C270 was targeted by GSSG to increase the activity of the protease, this unique residue might be associated with the severe consequences of SARS-CoV and SARS-CoV-2 infections to human beings. Additionally, the unique C270 of the SARS-CoV and SARS-CoV-2 PL^pro^ provides the structural basis for discovery of selective allosteric modulators.

## Conclusions

The pandemic of COVID-19 together with the epidemics of SARS and MERS has raised great awareness about the increasing infection risks of highly pathogenic CoVs, calling for a huge demand in discovery of anti-coronaviral drugs. PL^pro^ is a highly conserved cysteine proteinase that is indispensable for coronavirus replication, providing an attractive but challenging target for developing broad-spectrum antiviral drugs. Herein, we find a novel allosteric inhibitor of the SARS-CoV-2 PL^pro^ with a covalent action to C270 which is a previously unrecognized site for allosteric regulation of the protease’s activity. The mechanism of action of such a new inhibitor and structural determinants associated with its binding uniquely suited for engaging the non-catalytic cysteine are significantly distinctive to known inhibitors which bind to the catalytic site of the SARS-CoV-2 PL^pro^. Rather unexpectedly, the discovery of this allosteric site is derived from the identification of two activators, dimesna and pyritinol, from a high-throughput screening. Moreover, this vital site is also covalently modified by GSSG, an endogenous activator, which may link to intracellular proteolytic activity of the SARS-CoV-2 or SARS-CoV PL^pro^. Our study thereby highlights a promising strategy for identification of the unrecognized allosteric modulation site based on the discovery of covalent probes and reveals an exquisite allosteric modulation mechanism for the protease, which offers a great opportunity for the development of new inhibitors of the SARS-CoV PL^pro^.

## Acknowledgments

This work was supported by the National Natural Science Foundation of China (No. 21877122 and No. 32071248), the Science and Technology Commission of Shanghai Municipality (No. 20430780300 and No. TM202101H003), and the Qiusuo Outstanding Youth Project of Lingang Laboratory (No. LG-QS-202205-02).

## Supporting Information

## Experimental Procedures

### Chemicals

Dimesna (CAS No. 16208-51-8) and Pyritinol (CAS No. 1098-97-1) were purchased from Bide Pharmatech. Bis[2-(N,N-dimethylamino)ethyl] disulfide (DMGA, CAS No. 17339-60-5) was purchased from Jiuding Chemical Technology. All other chemicals, unless otherwise noted, were purchased from Sigma. The purity of all compounds used for biochemical test is over 95%.

### Plasmids construction

The SARS-CoV-2 papain-like protease (PL^pro^) protein sequence (pp1ab amino acids, 1564−1878) was obtained from the SARS-CoV-2 genome (National Center for Biotechnology Information Genome databank: NC_045512.2). For bacterial expression, the cDNA encoded the SARS-CoV-2 PL^pro^ with *E. coli* codon optimization was ordered from GenScript and cloned into the pET15b expression vector with an N-terminal 6 × His-SUMO2 fusion tag. For transfection of mammalian cell, the cDNA encoded the SARS-CoV-2 PL^pro^ with mammalian codon optimization was also ordered from GenScript and cloned into the pcDNA 3.1 with an C-terminal FLAG tag. The sequence of pcDNA3-PL-flipGFP-T2A-mCherry was designed based on plasmid pcDNA3-TEV-flipGFP-T2A-mCherry (Addgene catalog NO.124429) where TEV cleave site was replaced by SARS-CoV-2 PL^pro^ cleavage site (LRGGAPTK), and ordered from GenScript. The mutants used in this study were constructed by site mutation PCR and verified with sequencing.

### Protein expression and purification

The plasmids of SARS-CoV-2 PL^pro^ and its variants for protein expression were transformed into BL21 (DE3) cells. The expressed proteins were purified by a Ni-NTA column (GE) and cleaved by SUMO Specific Peptidase 2 (SENP2) for removing the 6 × His-SUMO2 fusion tag. The resulting protein samples were further purified by SP-Sepharose (GE Healthcare) and Superdex200 (GE Healthcare). The eluted proteins were stored in a solution containing 25 mM Tris pH7.5 and 2 mM DTT for the subsequent experiments.

### Enzymatic assay of the SARS-CoV-2 PL^pro^ or its variants

The enzymatic assay of all compounds were performed using 96-well plates. The substrate used in the assay was the fluorogeneic peptide (RLRGG-AMC)^[42]^, which was synthesized by GenScript. Reactions were performed in a total volume of 120 μL, which contained the following components: 50 mM Tris pH7.5, 0.1 mg/mL BSA, 50 nM SARS-CoV-2 PL^pro^ or its variants, indicated concentrations of compound or equal volume of solvent (DMSO or H_2_O). After incubation for 30 min, reactions were initiated with the addition of RLRGG-AMC to reach a final concentration of 20 µM. After that, the fluorescent signal was immediately measured every 1 min for 5 min with a BioTek H1 plate reader (excitation: 360 nm, emission: 460 nm). The initial velocity of reaction added with the compound at various concentrations compared to the reaction added with the solvent were calculated. This enzymatic assay was used to screen a library of 4210 compounds including the FDA-approved drugs and candidate compounds in clinical trials. In addition, this assay was also used to produce does-response curves of activators and inhibitors. Half maximal effective/inhibitory concentration (EC_50_ for activators, IC_50_ for inhibitors) and maximal effect (E_max_) were determined by nonlinear regression analysis of the does-response curves using GraphPad Prism.

### Determination of *K*_m_ and *V*_max_

The determination of Michaelis constant (*K*_m_) and maximum rate achieved by the enzyme (*V*_max_) were based on the enzymatic assay described above. Reactions were also performed in a total volume of 120 μL, containing 50 mM Tris pH 7.5, 0.1 mg/mL BSA, 50 nM SARS-CoV-2 PL^pro^ or its variants, and indicated concentrations of compounds or equal volume of solvent. After incubation for 30 min, reactions were initiated with the addition of various concentrations of the fluorogeneic substrate (RLRGG-AMC, 13.0-1666.7 μM). Initial velocities were measured as mentioned above and the velocity versus substrate concentration plot were further analyzed by Michaelis-Menten equation to obtain *K*_m_ and *V*_max_ using GraphPad Prism.

### Kinetic analysis of compounds covalently binding to the SARS-CoV-2 PL^pro^

The interaction of covalent compounds with the SARS-CoV-2 PL^pro^ can be described in two steps: an initial reversible binding event (*K*_i_ for inhibitors, *K*_a_ for activators) followed by formation of the covalent bond (*k*_incat_). For determination of *K*_i_ or *K*_a_ and *k*_inact_, 50 nM recombinant SARS-CoV-2 PL^pro^ was incubated with various concentrations of the compound for indicated time. At each time point, the enzymatic assay was carried out as mentioned above. Relative enzymatic activity was calculated by the initial velocity ratio of reactions added with compound over to the reaction added with solvent. For activators, the relative enzymatic activity needs to be normalized ((enzymatic-activity%-100%)/(E_max_-100%)). Relative enzymatic activity for various concentrations of the compound over a time course were fit to the semi-logarithmic plot equation to generate observed the rate constant value (*k*_obs_) for each concentration tested. The resulting *k*_obs_ values were then plotted versus compound concentrations ([C]), then *k*_inact_ and *K*_i_ or *K*_a_ values were calculated according to the equation: *k*_obs_ = *k*_inact_ × ([C]/([C] + *K*_i_ or *K*_a_)) using GraphPad Prism.

### Cell-based PL-FlipGFP assay

This assay was designed stemming from previous reports^[31, 52]^ In detail, a FlipGFP in which the 10th and 11th β-strands were separated from the rest of the GFP β-barrel (β-strands 1-9) were used. The 10th β-strand, the coiled-coil E5, the 11th β-strand, the cleavage sequence (LRGGAPTK) by the SARS-CoV-2 PL^pro^, and the coiled-coil K5 were linked in order (Figure S2). In the absence of the SARS-CoV-2 PL^pro^, the FlipGFP is inactive, as the 10th and 11th β-strands were restrained due to the presence of heterodimerized coiled-coil E5/K5 and unable to associate with the rest of GFP β-barrel. Upon the cleavage by the protease, the 10th or 11th β-strands then flips to associate with β-strands 1-9, leading to restoration of the integrated GFP for green fluorescence detection. Meanwhile, a red fluorescent protein mCherry was included via a “self-cleaving” 2A peptide to act as an internal control, and the ratio of green over red fluorescent signal (GFP/mCherry) represents the amount of products generated by the protease hydrolysis in cells.

In the present study, HEK293T cells were seeded in 12-well plates overnight. The following day, cells at 70-90% confluency were co-transfected with 500 ng of pcDNA3-PL-flipGFP-T2A-mCherry plasmid and various amounts (20-100 ng) of protease expression plasmids in each well with 2.4 μg polyethylenimine (PEI). After 48 hours, the supernatant was removed and 200 μL lysis buffer (100 mM Tris pH7.5, 100 mM NaCl, 1 mM EDTA, 1% Triton X-100) was added. After incubation for 5 min at 4 °C, 120 μL cell lysate was transferred to a 96-well plate and the fluorescent signal of GFP (excitation: 475 nm, emission: 505 nm) and mCherry (excitation: 580 nm, emission: 610 nm) were measured. The amount of hydrolyzed product in cells was calculated based on the ratio of GFP/mCherry fluorescent signal. After that, these samples were denatured with 4x Laemmli buffer added with β-mercaptoethanol and heated at 95 °C for 10 min for the subsequent immunoblotting test.

### S-glutathionylation of the SARS-CoV-2 PL^pro^

Concentrated protease (the wild-type or C270S variant of SARS-CoV-2 PL^pro^) was diluted with 50 mM Tris pH7.5 to a final concentration of 0.5 mg/mL. 50 μL diluted protease were then incubated with 5 mM GSSG or equivalent volume of solvent at 25 °C for 30 min. After that, the samples were divided into two parts. Under the reduced condition, samples were additionally added with 1% β-mercaptoethanol. In the end, all samples were denatured with 4x Laemmli buffer (without any reduced agents) containing 1 mM N-ethyl maleimide for the alkylation of accessible thiols.

### Immunoblotting

All denatured samples (10 μL) were run on the 12.5% acrylamide SDS-PAGE gels. After that, proteins were transferred to 0.2 μm polyvinylidene difluoride membranes (Bio-Rad, 160177). Membranes were washed with TBST (50 mM Tris pH7.5, 150 mM NaCl, 0.1% Tween 20) and blocked with 5% bovine serum albumin (BSA) in TBST for 1 h at 25 °C. Membranes were first incubated with the primary antibody (Anti-Glutathione antibody [D8], ab19534, Abcam; Mouse anti DDDDK-Tag (FLAG-tag) mAb, AE005, ABclonal) in 5% BSA TBST for 16 h at 4 °C, washed with TBST three times, then incubated with the secondary antibody (HRP-labeled Goat Anti-Mouse IgG(H+L), A0216, Beyotime) in 5% BSA TBST (1 h, 25 °C), and washed with TBST three times. Imaging solution (Tanon High-sig ECL Western Blotting Substrate, 180-501) was added to the membranes, and chemiluminescence was detected on the Tanon 5200CE using standard chemiluminescence settings.

## Supplementary Figures

**Figure S1.**
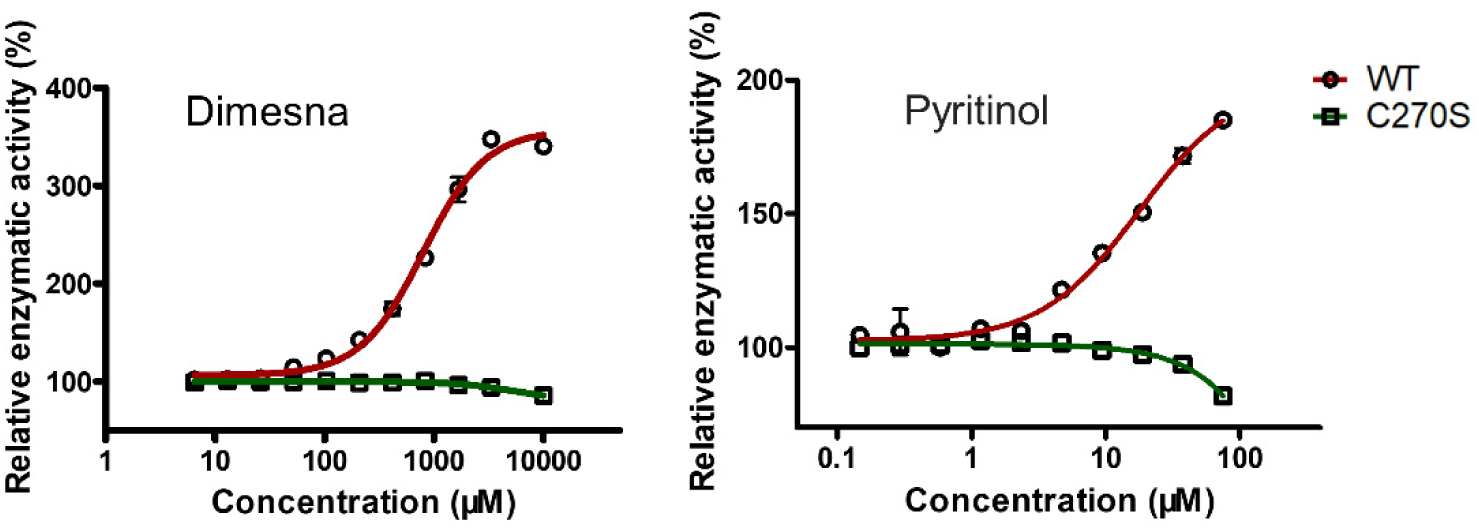
Representative profiles for the activation of dimesna (left panel) and pyritinol (right panel) on the wild-type (red) and C270S variant (green) of SARS-CoV-2 PL^pro^.

**Figure S2.**
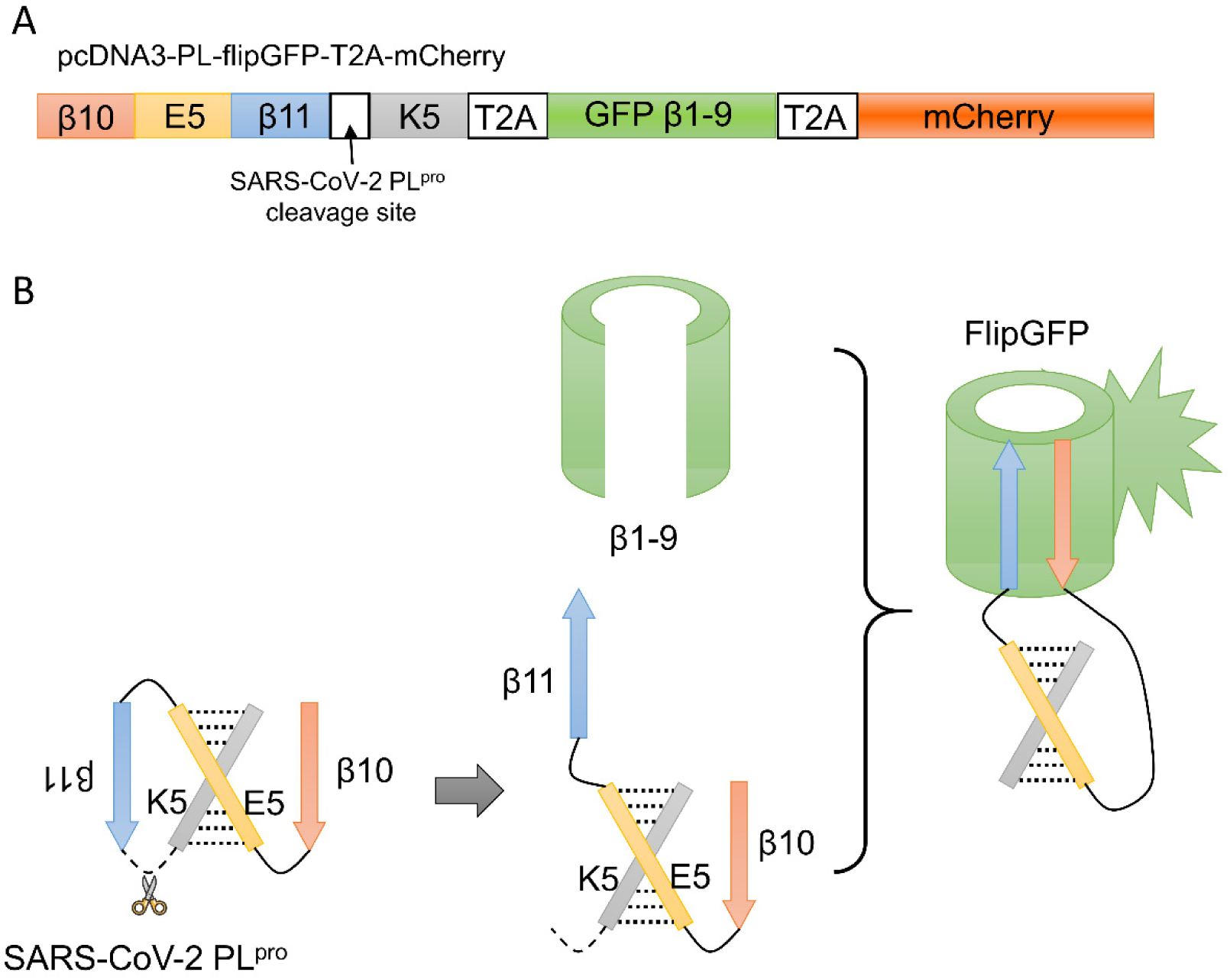
Schematic diagram of pcDNA3-PL-flipGFP-T2A-mCherry (A) and the cell-based FlipGFP assay (B).

**Figure S3.**
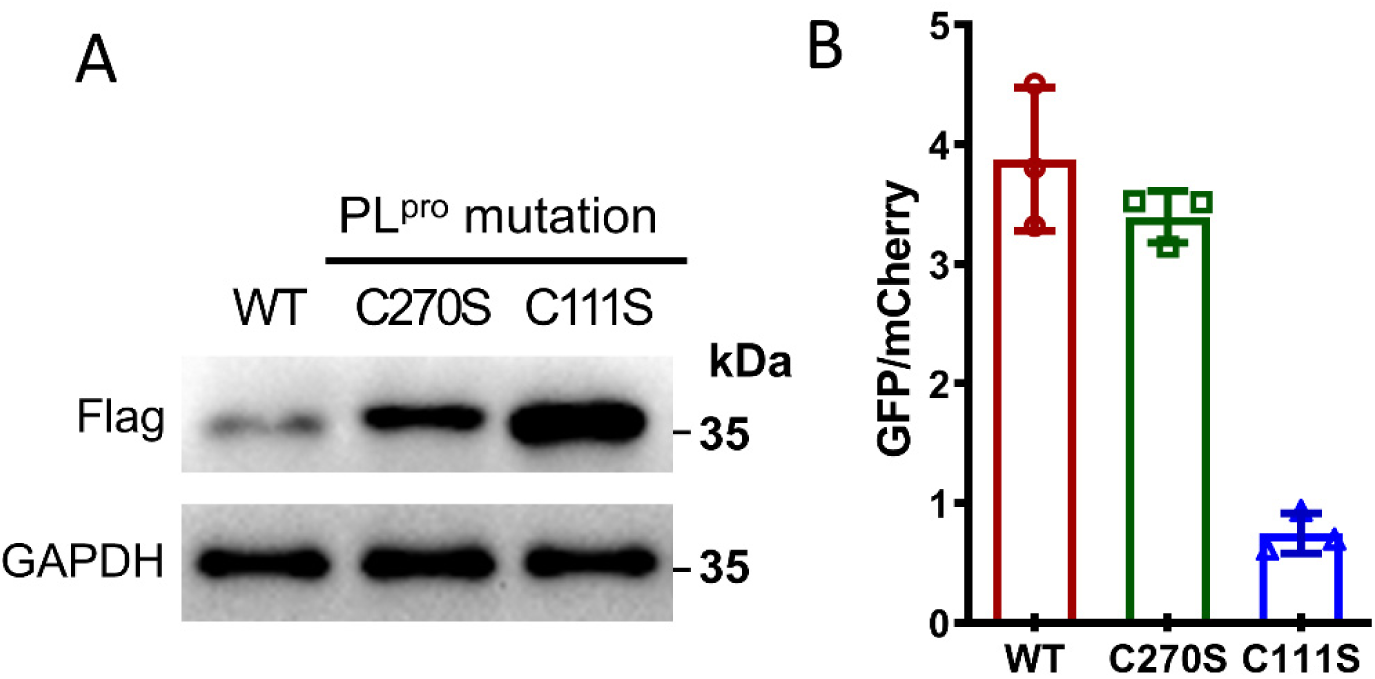
Protein expression level (A) and in-cell activity (B) of the wide-type (WT) or C270S variant of SARS-CoV-2 PL^pro^ after transfection with same amount of plasmids.

**Figure S4.**
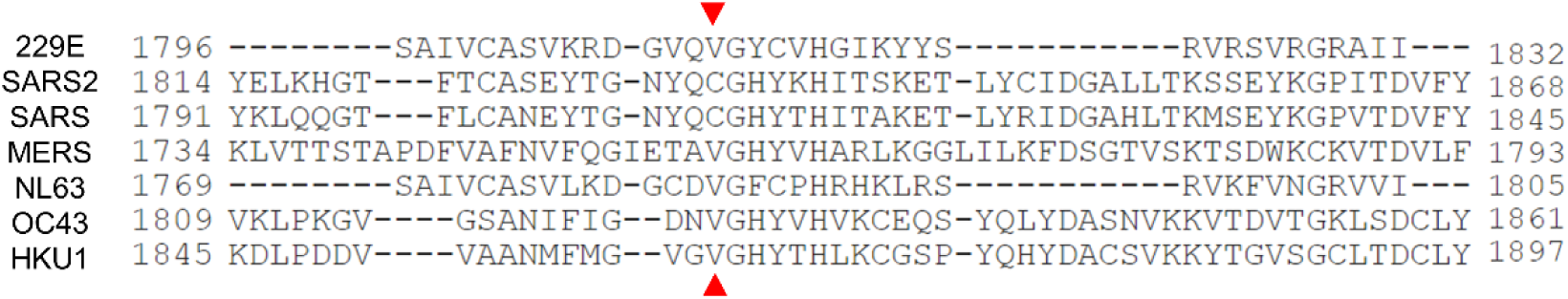
Multiple alignment of seven human COVs’ pp1ab protein sequences generated by Clustal Omega program^[53]^ (https://www.uniprot.org/align/). Red triangle marks the equivalent position of C270 on SARS-CoV-2 PL^pro^. Only the window of the amino acids nearby C270 was shown for clarity reasons.

## References

[1] Zumla A, Chan J F, Azhar E I, et al. Coronaviruses - drug discovery and therapeutic options[J]. Nat. Rev. Drug Discov.: 2016, 15: 327–347.

[2] de Wit E, van Doremalen N, Falzarano D, et al. SARS and MERS: recent insights into emerging coronaviruses[J]. Nat. Rev. Microbiol.: 2016, 14: 523–534.

[3] Chan J F, Yuan S, Kok K H, et al. A familial cluster of pneumonia associated with the 2019 novel coronavirus indicating person-to-person transmission: a study of a family cluster[J]. Lancet: 2020, 395: 514–523.

[4] Li Q, Guan X, Wu P, et al. Early transmission dynamics in Wuhan, China, of novel coronavirus-infected pneumonia[J]. N. Engl. J. Med.: 2020, 382: 1199–1207.

[5] Cui J, Li F, Shi Z L. Origin and evolution of pathogenic coronaviruses[J]. Nat. Rev. Microbiol.: 2019, 17: 181–192.

[6] Su S, Wong G, Shi W, et al. Epidemiology, genetic recombination, and pathogenesis of coronaviruses[J]. Trends Microbiol.: 2016, 24: 490–502.

[7] Zhou P, Yang X L, Wang X G, et al. A pneumonia outbreak associated with a new coronavirus of probable bat origin[J]. Nature: 2020, 579: 270–273.

[8] Hajjar L A, Costa I B S D, Rizk S I, et al. Intensive care management of patients with COVID-19: a practical approach[J]. Annals of Intensive Care: 2021, 11: 36.

[9] Harrison A G, Lin T, Wang P. Mechanisms of SARS-CoV-2 transmission and pathogenesis[J]. Trends Immunol.: 2020, 41: 1100–1115.

[10] V’Kovski P, Kratzel A, Steiner S, et al. Coronavirus biology and replication: implications for SARS-CoV-2[J]. Nat. Rev. Microbiol.: 2021, 19: 155–170.

[11] Xiong M Y, Su H X, Zhao W F, et al. What coronavirus 3C-like protease tells us: From structure, substrate selectivity, to inhibitor design[J]. Med. Res. Rev.: 2021, 41: 1965–1998.

[12] Lei J, Kusov Y, Hilgenfeld R. Nsp3 of coronaviruses: Structures and functions of a large multi-domain protein[J]. Antiviral Res.: 2018, 149: 58–74.

[13] Klemm T, Ebert G, Calleja D J, et al. Mechanism and inhibition of the papain-like protease, PLpro, of SARS-CoV-2[J]. EMBO J.: 2020, 39: e106275.

[14] Shin D, Mukherjee R, Grewe D, et al. Papain-like protease regulates SARS-CoV-2 viral spread and innate immunity[J]. Nature: 2020, 587: 657–662.

[15] Ratia K, Kilianski A, Baez-Santos Y M, et al. Structural basis for the ubiquitin-linkage specificity and deISGylating activity of SARS-CoV papain-like protease[J]. PLoS Pathog.: 2014, 10: e1004113.

[16] Barretto N, Jukneliene D, Ratia K, et al. The papain-like protease of severe acute respiratory syndrome coronavirus has deubiquitinating activity[J]. J. Virol.: 2005, 79: 15189–15198.

[17] Freitas B T, Durie I A, Murray J, et al. Characterization and noncovalent inhibition of the deubiquitinase and deISGylase activity of SARS-CoV-2 papain-like protease[J]. ACS Infect. Dis.: 2020, 6: 2099–2109.

[18] Chen X J, Yang X X, Zheng Y, et al. SARS coronavirus papain-like protease inhibits the type I interferon signaling pathway through interaction with the STING-TRAF3-TBK1 complex[J]. Protein Cell: 2014, 5: 369–381.

[19] Frieman M, Ratia K, Johnston R E, et al. Severe acute respiratory syndrome coronavirus papain-like protease ubiquitin-like domain and catalytic domain regulate antagonism of IRF3 and NF-kappa B signaling[J]. J. Virol.: 2009, 83: 6689–6705.

[20] Liu G Q, Lee J H, Parker Z M, et al. ISG15-dependent activation of the sensor MDA5 is antagonized by the SARS-CoV-2 papain-like protease to evade host innate immunity[J]. Nat. Microbiol.: 2021, 6: 467–478.

[21] Munnur D, Teo Q W, Eggermont D, et al. Altered ISGylation drives aberrant macrophage-dependent immune responses during SARS-CoV-2 infection[J]. Nat. Immunol.: 2021, 22: 1416–1427.

[22] Del Valle D M, Kim-Schulze S, Huang H H, et al. An inflammatory cytokine signature predicts COVID-19 severity and survival[J]. Nat. Med.: 2020, 26: 1636–1634.

[23] Qiao J, Li Y S, Zeng R, et al. SARS-CoV-2 M(pro) inhibitors with antiviral activity in a transgenic mouse model[J]. Science: 2021, 371: 1374–1378.

[24] Zhang L, Lin D, Sun X, et al. Crystal structure of SARS-CoV-2 main protease provides a basis for design of improved alpha-ketoamide inhibitors[J]. Science: 2020, 368: 409–412.

[25] Dai W, Zhang B, Jiang X M, et al. Structure-based design of antiviral drug candidates targeting the SARS-CoV-2 main protease[J]. Science: 2020, 368: 1331–1335.

[26] Su H X, Yao S, Zhao W F, et al. Anti-SARS-CoV-2 activities in vitro of Shuanghuanglian preparations and bioactive ingredients[J]. Acta Pharmacol. Sin.: 2020, 41: 1167–1177.

[27] Su H, Yao S, Zhao W, et al. Identification of pyrogallol as a warhead in design of covalent inhibitors for the SARS-CoV-2 3CL protease[J]. Nat. Commun.: 2021, 12: 3623.

[28] Owen D R, Allerton C M N, Anderson A S, et al. An oral SARS-CoV-2 M(pro) inhibitor clinical candidate for the treatment of COVID-19[J]. Science: 2021, 374: 1586–1593.

[29] Boras B, Jones R M, Anson B J, et al. Preclinical characterization of an intravenous coronavirus 3CL protease inhibitor for the potential treatment of COVID19[J]. Nat. Commun.: 2021, 12: 6055.

[30] Shen Z, Ratia K, Cooper L, et al. Design of SARS-CoV-2 PLpro inhibitors for COVID-19 antiviral therapy leveraging binding cooperativity[J]. J. Med. Chem.: 2022, 65: 2940–2955.

[31] Ma C, Sacco M D, Xia Z, et al. Discovery of SARS-CoV-2 papain-like protease inhibitors through a combination of high-throughput screening and a FlipGFP-based reporter assay[J]. ACS Cent. Sci.: 2021, 7: 1245–1260.

[32] Ghosh A K, Brindisi M, Shahabi D, et al. Drug development and medicinal chemistry efforts toward SARS-coronavirus and Covid-19 therapeutics[J]. Chemmedchem: 2020, 15: 907–932.

[33] Rut W, Lv Z, Zmudzinski M, et al. Activity profiling and crystal structures of inhibitor-bound SARS-CoV-2 papain-like protease: A framework for anti-COVID-19 drug design[J]. Sci. Adv.: 2020, 6: eabd4596.

[34] Cho C C, Li S G, Lalonde T J, et al. Drug repurposing for the SARS-CoV-2 papain-like protease[J]. Chemmedchem: 2022, 17: e202100455.

[35] Shan H, Liu J, Shen J, et al. Development of potent and selective inhibitors targeting the papain-like protease of SARS-CoV-2[J]. Cell Chem. Biol.: 2021, 28: 855–865.

[36] Zhao Y, Du X Y, Duan Y K, et al. High-throughput screening identifies established drugs as SARS-CoV-2 PLpro inhibitors[J]. Protein Cell: 2021, 12: 877–888.

[37] Ratia K, Pegan S, Takayama J, et al. A noncovalent class of papain-like protease/deubiquitinase inhibitors blocks SARS virus replication[J]. Proc. Natl. Acad. Sci. U. S. A.: 2008, 105: 16119–16124.

[38] Osipiuk J, Azizi S A, Dvorkin S, et al. Structure of papain-like protease from SARS-CoV-2 and its complexes with non-covalent inhibitors[J]. Nat. Commun.: 2021, 12: 743.

[39] Fu Z Y, Huang B, Tang J L, et al. The complex structure of GRL0617 and SARS-CoV-2 PLpro reveals a hot spot for antiviral drug discovery[J]. Nat. Commun.: 2021, 12.

[40] Shan H Y, Liu J P, Shen J L, et al. Development of potent and selective inhibitors targeting the papain-like protease of SARS-CoV-2[J]. Cell Chem. Biol.: 2021, 28: 855–865.

[41] Gao X, Qin B, Chen P, et al. Crystal structure of SARS-CoV-2 papain-like protease[J]. Acta Pharm. Sin. B: 2021, 11: 237–245.

[42] Ghosh A K, Takayama J, Aubin Y, et al. Structure-based design, synthesis, and biological evaluation of a series of novel and reversible inhibitors for the severe acute respiratory syndrome-coronavirus papain-like protease[J]. J. Med. Chem.: 2009, 52: 5228–5240.

[43] Sadowsky J D, Burlingame M A, Wolan D W, et al. Turning a protein kinase on or off from a single allosteric site via disulfide trapping[J]. Proc. Natl. Acad. Sci. U. S. A.: 2011, 108: 6056–6061.

[44] Nussinov R, Tsai C J. The design of covalent allosteric drugs[J]. Annu. Rev. Pharmacol. Toxicol.: 2015, 55: 249–267.

[45] Darby J F, Atobe M, Firth J D, et al. Increase of enzyme activity through specific covalent modification with fragments[J]. Chem. Sci.: 2017, 8: 7772–7779.

[46] Ostrem J M, Peters U, Sos M L, et al. K-Ras(G12C) inhibitors allosterically control GTP affinity and effector interactions[J]. Nature: 2013, 503: 548–551.

[47] Hardy J A, Wells J A. Searching for new allosteric sites in enzymes[J]. Curr. Opin. Struct. Biol.: 2004, 14: 706–715.

[48] Erlanson D A, Wells J A, Braisted A C. Tethering: fragment-based drug discovery[J]. Annu. Rev. Biophys. Biomol. Struct.: 2004, 33: 199–223.

[49] Henderson J A, Verma N, Harris R C, et al. Assessment of proton-coupled conformational dynamics of SARS and MERS coronavirus papain-like proteases: Implication for designing broad-spectrum antiviral inhibitors[J]. J. Chem. Phys.: 2020, 153: 115101.

[50] Hwang C, Sinskey A J, Lodish H F. Oxidized redox state of glutathione in the endoplasmic reticulum[J]. Science: 1992, 257: 1496–1502.

[51] Montero D, Tachibana C, Rahr Winther J, et al. Intracellular glutathione pools are heterogeneously concentrated[J]. Redox Biol.: 2013, 1: 508–513.

[52] Q. Zhang, A. Schepis, H. Huang, J. Yang, W. Ma, J. Torra, S. Q. Zhang, L. Yang, H. Wu, S. Nonell, Z. Dong, T. B. Kornberg, S. R. Coughlin, X. Shu, J. Am. Chem. Soc.: 2019, 141: 4526–4530.

[53] J. Soding, Bioinformatics: 2005, 21: 951–960.

